# Short-term heat stress assays provide a standardized screening tool to assess impacts of microbiome manipulation on Aiptasia thermal stress tolerance

**DOI:** 10.1101/2023.06.01.543236

**Authors:** Melanie Dörr, Julia Denger, Céline S. Maier, Jana V. Kirsch, Hannah Manns, Christian R. Voolstra

**Author notes:** These authors contributed equally to this work (Julia Denger, Céline S. Maier).

## Abstract

The ongoing loss of corals and their reef ecosystems hastens the need to develop approaches that mitigate the impacts of climate change. Given the strong reliance of corals on their associated prokaryotic and microalgal symbionts, microbiome-targeted interventions in the form of probiotics or microbiome transplants are emerging as potential solutions. Although inoculation with beneficial microorganisms was shown to improve coral bleaching recovery, the mechanistic underpinnings and extent to which microbiomes can be manipulated are largely unknown. Research progress in this regard is often hindered by coral holobiont complexity and a lack of standardized diagnostics to assess physiological and phenotypic changes following microbial manipulation. Here we address these shortcomings by establishing short-term acute thermal stress assays using the CBASS (Coral Bleaching Automated Stress System) as a standardized and reproducible experimental platform to assess the impact of microbiome manipulation on stress tolerance phenotypes of the coral model Aiptasia. We show that thermal tolerance phenotypes following acute heat stress assays are highly reproducible, host species-specific, and can exert legacy effects with consequences for long-term thermal resilience. We further demonstrate the ability to resolve phenotypic differences in thermal tolerance following microbiome manipulation through pathogen incubation. By employing acute heat stress assays in conjunction with a tractable model organism, we present a standardized experimental platform that allows functional screening for microbes that affect thermal stress tolerance. Such effort may accelerate the discovery of microbes and microbial mechanisms mediating thermal tolerance and our ability to harness them to increase stress resilience.

## Introduction

Recent records indicate that reef ecosystem decline is accelerating at a pace quickly surpassing the adaptability of coral individuals (Bennett et al., 2021). Anthropogenic climate change and resulting ocean warming lead to global coral bleaching events with increasing frequency and severity (Hughes et al., 2018b). Current estimates suggest that 98% of the Great Barrier Reef has been subjected to bleaching at least once following widespread mass bleaching events in 2016, 2017, and, most recently, 2020 (Hughes et al., 2021; Eakin et al., 2022). This is just one example demonstrating how coral bleaching, i.e., the loss of dinoflagellate algal symbionts (family Symbiodiniaceae) (LaJeunesse et al., 2018), has become one of the main drivers of coral mortality and reef degradation (Hughes et al., 2018a; Suggett and Smith, 2020; Eakin et al., 2022). Future projections are grim, estimating that a quarter of all reefs have already been lost due to climate impacts, and the overwhelming majority is at risk of extinction if global warming is not curbed to below 2 °C in the coming decades (Allen et al., 2018). Thus, there is an urgent need to devise and implement active interventions to support restoration and conservation efforts (Kleypas et al., 2021; Knowlton et al., 2021; Voolstra et al., 2021a, 2023).

Corals are not only dependent on their algal symbionts but also form strong associations with an array of other microorganisms contributing to host physiology (Rosenberg et al., 2007). In particular, the bacterial microbiome, i.e., the sum of all associated bacteria, is more variable than algal associations and commonly adjusts to and reflects prevailing environmental conditions (Reshef et al., 2006; Roder et al., 2015; Ziegler et al., 2019), with evidence that changes in the microbiome support environmental adaptation, e.g., to counter thermal stress (Ziegler et al., 2017). Consequently, microbiome-targeted interventions in the form of probiotics or microbiome transplants are emerging as potential solutions to support coral resilience (Doering et al., 2021; Santoro et al., 2021; Peixoto et al., 2022). To this end, proof-of-principle of probiotic treatment during heat stress demonstrated an increased recovery from bleaching and coral survival (Rosado et al., 2019; Santoro et al., 2021). However, we are still far away from a mechanistic understanding of how bacteria increase holobiont resilience and the extent to which microbiomes can be manipulated, given that not all species exhibit ‘microbiome flexibility’ (Ziegler et al., 2019; Voolstra and Ziegler, 2020), or how long such effects remain.

The development of standardized approaches can greatly facilitate study comparison and advance insight (Grottoli et al., 2021). Platforms such as the Coral Bleaching Automated Stress System (CBASS), for instance, have been shown to promptly resolve thermotolerance differences across species and sites with subsequent elucidation of holobiont differences and insights into the underlying mechanisms (Voolstra et al., 2020, 2021b; Savary et al., 2021). More recently, CBASS assays were used to assess the impact of lowered oxygen levels on bleaching susceptibility by comparing standardized thermal tolerance thresholds, i.e., ED50s (Evensen et al., 2021, 2022), of corals under normative and hypoxic O_2_ seawater levels (Alderdice et al., 2022a). Thus, CBASS assays cannot only be employed to determine coral *in situ* thermal tolerance thresholds but also to assess how those thresholds change under the impact of other factors.

Here, we sought to test the efficacy of CBASS assays as a standardized screening tool to assess the impact of microbiome manipulation on thermal tolerance phenotypes using the coral model Aiptasia (Baumgarten et al., 2015; Costa et al., 2021). We show that (i) CBASS assays are highly reproducible, (ii) that repeated heat stress exposures exhibit legacy effects consequential to thermal tolerance measurements and long-term resilience, and (iii) that CBASS can reliably resolve differences in thermal tolerance following microbiome manipulation (i.e., incubation with the bleaching pathogen *Vibrio coralliilyticus*). Our data support the use of CBASS assays as a standardized experimental platform for the screening and functional interrogation of microbes that affect thermal stress tolerance through identification of altered bleaching phenotypes (i.e., ED50 thermal tolerance thresholds) following exposure to defined bacterial isolates. Such effort should help to increase our understanding of the contribution of microbes to coral resilience and our ability to harness them.

## Materials & Methods

### Aiptasia rearing conditions

Clonal symbiotic Aiptasia (*sensu Exaiptasia diaphana*) (Dungan et al., 2020) anemones from the strains F003 (Carolina Biological Supply Company; 162865) (Grawunder et al., 2015) and H2 (Xiang et al., 2013) were maintained in translucent polycarbonate tanks filled with artificial seawater (ASW; PRO-REEF Sea Salt, Tropic Marin) at 35 g/L salinity. Animals were kept at a 25 °C diurnal 12 h light: 12 h dark cycle (12L:12D) under white LED lights with an intensity of ∼ 70 - 80 μmol m^−2^s^−1^ of photosynthetic active radiation (PAR). Animals were fed once a week with freshly hatched *Artemia* nauplii (Ocean Nutrition). Tanks were cleaned and supplied with fresh ASW the next day.

### Short-term acute heat stress assays (CBASS)

Short-term acute heat stress assays on Aiptasia anemones were performed using the Coral Bleaching Automated Stress System (CBASS). The CBASS setup was based on (Voolstra et al., 2020) using four 10 L tanks per system (Figure S1), which allows to simultaneously run four independent temperature profiles (Figure S2). Temperature profiles were controlled using programmable thermostats (Inkbird ITC-310T-B). The temperature in each tank was adjusted by two IceProbe Thermoelectric chillers (Nova Tec) and one 200W titanium aquarium heater (Schego) that were connected to the Inkbird thermostat. HOBO Pendant Temperature Loggers were used to log temperature profiles of each tank in 5 min intervals (Figure S3). A dimmable 165 W full spectrum LED aquarium light (Galaxyhydro) was used for each tank. Light settings were adjusted to ∼ 80 μmol photons m^-2^s^-1^ to match long-term Aiptasia culturing conditions using an MQ-150 Underwater Full Spectrum Quantum Sensor (Apogee). For each CBASS run, anemones were placed into the respective tanks at 12.30 h, and temperature profiles were started at 13.00 h (Figure S3). The temperature in the control tank was maintained at 30 °C for the duration of the CBASS assay, i.e., 18 hours. The temperatures in the three heat stress treatment tanks were incrementally ramped up over 3 hours to 34 °C (control +4 °C, medium), 36 °C (control +6 °C, high) and 39 °C (control +9 °C, extreme). The heat stress temperatures were then held for 3 hours before ramping back down to 30 °C within 1 h (19.00 h), followed by an 11-h overnight recovery period until the following morning. In addition, lights were turned off after the heat-hold. Lights were turned back on between 7.00 h and 8.00 h, completing the 18 h short-term acute heat stress assay.

### Photosynthetic efficiency measurements and ED50 thermal tolerance thresholds

Dark-acclimated photosynthetic efficiency (F_v_/F_m_) of photosystem II of each anemone across all temperature treatments was measured after the ramp down to 30 °C (19.00 h) and following 1 h of dark acclimation (20.00 h). Measurements were also taken in the morning (7.00 h) at the end of the overnight baseline temperature hold (recovery period) before the lights were turned back on (Figure S4). The photosynthetic efficiency was measured using a MINI-PAM-II fluorometer (Walz, Germany). Temperature tolerance thresholds were determined based on Pulse Amplitude Modulation (PAM) fluorometry measurements (Voolstra et al., 2020). Given the clonal nature of Aiptasia anemones, temperature tolerance thresholds were determined per cohort of Aiptasia anemones (i.e., across all anemone replicates rather than individually) as the temperature at which the photosynthetic efficiency declined to 50% of the baseline temperature value. We refer to this as the Population Effective Dose 50 (ED50) that was computed using the DRC package in R (R4.1.2) (Ritz et al., 2015; Evensen et al., 2021). The standard error of population ED50s was extracted from the drm model using the ED() function.

### Reproducibility of Aiptasia population thermal tolerance thresholds, impact of repeated heat stress cycling, and heat stress legacy effects

Two independent CBASS systems (system A and system B) were simultaneously run to test the reproducibility of experimentally determined ED50 thermal tolerance thresholds. Each of the four temperature treatment tanks per CBASS system (control, medium, high, and extreme temperature profiles) housed 10 replicate anemones of either F003 or H2, which were kept in 25 mL Eppendorf tubes filled with 20 mL filtered artificial seawater (FASW, PRO-REEF Sea Salt, Tropic Marin) (Figure S1). F003 and H2 anemones were run in independent CBASS systems A and B (total of 4 CBASS systems). To assess the impact of repeated CBASS thermal stress testing on ED50 thermal thresholds, Aiptasia anemones were subjected to two consecutive CBASS runs (day 1, day 2). To further assess the presence of putative heat stress legacy effects and their impact on ED50 thermal tolerance thresholds, surviving animals from the two consecutive runs were kept under rearing conditions for one month to allow for putative recovery and were subsequently subjected to another CBASS run. All anemones from the 39 °C heat stress treatment died during the one-month interval and their F_v_/F_m_ values were set to 0 for ED50 thermal tolerance thresholds modeling. For the remaining temperature treatments (i.e., 30 °C, 33 °C, 36 °C), surviving animals were available to measure F_v_/F_m_ following heat stress.

### Microbiome manipulation through bacteria inoculation

*Vibrio coralliilyticus* (isolated from blacktip reef sharks, d’Arros Island of Amirante Archipel, Seychelles) was revived from -80 °C by streaking on marine agar (37.4 g marine broth (Difco) and 17.7 g agar in 1 L deionized water) to check viability and purity (Pogoreutz et al., 2019). A single bacterial colony was inoculated in marine broth and incubated for 24 h at 25 °C to reach an optical density of approximately OD_600_ = 0.8 (equal to 10^8^-10^9^ CFU/mL) following a previous study (Ushijima et al., 2020), and measured using a CLARIOstar Plus Plate Reader (BMG Labtech). The culture was harvested at 3,000 x g for 5 min and resuspended in FASW. Prior to bacterial inoculation, the culture was diluted with FASW to a final concentration of 10^7^ cells/mL. Five anemones per strain and temperature were each inoculated with 20 mL FASW containing the liquid culture at 10^7^ cells/mL in 25 mL Eppendorf tubes 24 h prior to CBASS. Anemones were kept being exposed to *V. coralliilyticus* during the 18 h short-term acute heat stress assay. Temperature profiles were identical to those above (30 °C, 34 °C, 36 °C, 39 °C), and dark-acclimated photosynthetic efficiency was measured after the temperature ramp down and accompanying dark acclimation for one hour (see above).

## Results

### High reproducibility of CBASS-based thermal tolerance thresholds

To assess the reproducibility of ED50-based thermal tolerance thresholds, we ran CBASS assays of 10 anemones each of strains F003 and H2 in duplicate using two independent systems (system A and system B comprised each of 4 temperature profile tanks) (Figure 1 A). We found that anemones of a given strain exhibited highly reproducible thermal tolerance thresholds between replicate CBASS assays: based on ED50 values, we found a 0.13 °C difference between both F003 anemone runs and only a 0.02 °C difference between the replicate H2 anemone runs (Figure 1 B, Table S1). Moreover, the distribution of dark-acclimated photosynthetic efficiencies for both sets of anemones across the replicate CBASS runs was largely identical, as apparent from the ED50 regression curves and data distribution. In comparison to F003 anemones, H2 anemones showed slightly lower ED50 thermal thresholds.

**Figure 1.**
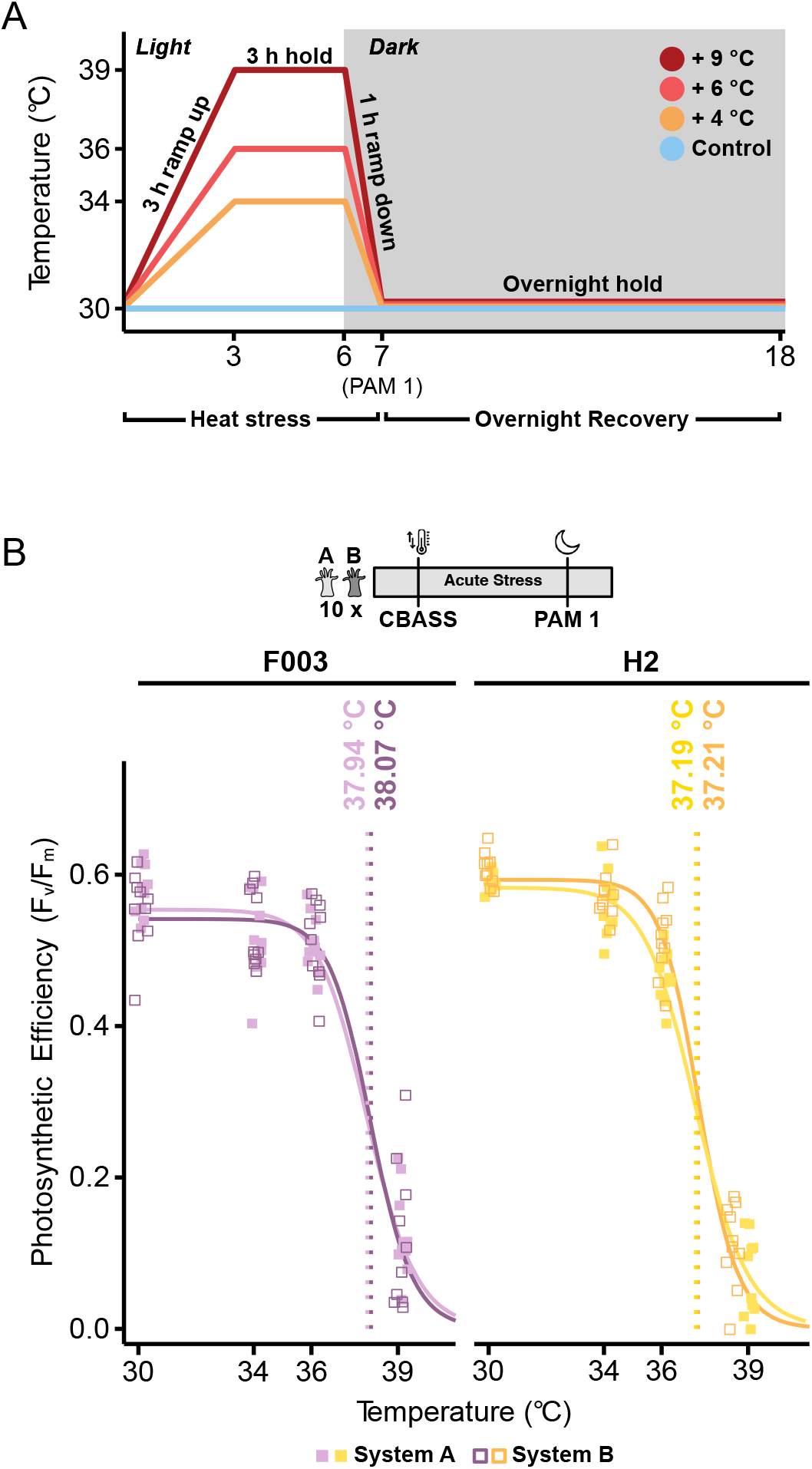
High reproducibility of CBASS thermal tolerance thresholds. Anemones in two separate CBASS systems show highly similar thermal tolerance thresholds. **(A)** Schematic representation of the short-term acute heat stress assay using the Coral Bleaching Automated Stress System (CBASS; adapted from Voolstra et al., 2020). The 18 h CBASS profile was run at 30 °C (control) and with a 3 h heat-hold at 34 °C (control +4 °C, medium), 36 °C (control +6 °C, high), and 39 °C (control +9 °C, extreme). Dark-acclimated photosynthetic efficiency was measured using a PAM fluorometer 1 h after temperature ramp down and accompanying dark acclimation (7 h, PAM 1). **(B)** Photosynthetic efficiencies (F_v_/F_m_) across replicated CBASS assays of Aiptasia strains F003 (dark and light purple) and H2 (yellow and orange) using two independent CBASS systems (system A and system B). Population ED50 thermal tolerance thresholds (n = 10 anemones per temperature profile) are denoted as vertical lines. Population ED50s are based on log-logistic regression curves of F_v_/F_m_ measurements across experimental temperatures and provide a standardized proxy for bleaching susceptibility (Evensen et al., 2021). ED50s and regression curves are largely identical and reproducible between the replicated CBASS runs.

### Repeated heat stress cycling affects CBASS-based thermal tolerance thresholds and exerts long-term legacy effects

To assess the impact of repeated heat stress cycles on CBASS-based thermal tolerance thresholds, we ran a repeated CBASS assay on the same set of 10 anemones per strain (F003 and H2) on consecutive days. To further assess the presence of long-term legacy effects that may arise from prior heat stress exposure, as alluded previously (Evensen et al., 2022), we subjected the same set of anemones to a third CBASS assay after one month of recovery (Figure 2 A). To follow photosynthetic efficiency over time, we measured dark-acclimated maximum quantum yields over three time points. Measurements were conducted after 1 hour of temperature ramp down and dark acclimation (according to the standardized CBASS protocol). We found that thermal tolerance thresholds (i.e., population ED50s) of F003 and H2 anemones were substantially lower in the second consecutive CBASS assay run (CBASS run 1 vs. CBASS run 2, Figure 2 B, C). F003 anemones showed a slightly larger loss of thermal tolerance than H2 anemones, in that the thermal tolerance difference between the first and second CBASS run was 1.77 °C for F003 and 1.61 °C for H2, although F003 exhibited overall higher thresholds than H2 (F003 ED50_CBASS1_ = 38.07 °C, ED50_CBASS2_ = 36.30 °C vs. H2 ED50_CBASS1_ = 37.21 °C, ED50_CBASS2_ = 35.60 °C) (Table S1).

**Figure 2.**
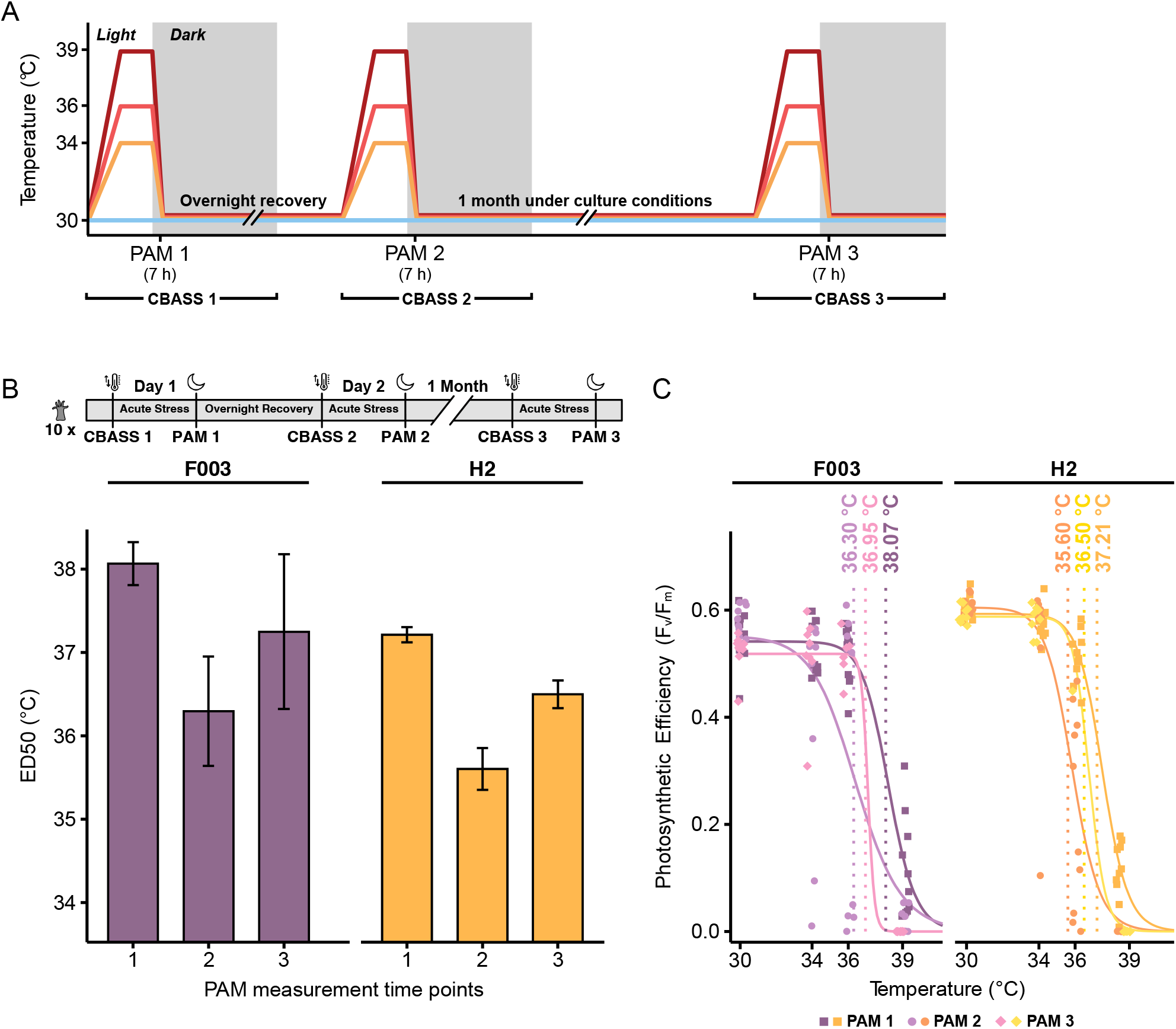
Repeated heat stress cycling decreases Aiptasia temperature tolerance thresholds and exerts long-term legacy effects. **(A)** Schematic representation of repeat CBASS runs. The same set of F003 and H2 anemones (n = 10 per temperature profile) were subjected to repeat CBASS runs on consecutive days (CBASS 1 and CBASS 2) as well as after one month under ambient rearing conditions (CBASS 3). PAM fluorometry measurement time points and CBASS cycles are indicated on the x-axis. **(B)** Population ED50 thermal tolerance thresholds based on photosynthetic efficiency (F_v_/F_m_) measures of Aiptasia strains F003 and H2 across three time points. ED50 thermal tolerance thresholds decrease with repeated stress testing but increase again after subsequent rearing under ambient conditions, although not to the initial level. Error bars represent the standard error of the population ED50s. **(C)** Photosynthetic efficiencies (F_v_/F_m_) across treatment temperatures for the three CBASS runs for F003 and H2 anemones. Please refer to (B) for an overview of the time points. Population ED50 thermal tolerance thresholds based on log-logistic regression curves of F_v_/F_m_ measurements are denoted as vertical lines.

To test the presence of putative long-term legacy effects following repeated exposure to heat stress (i.e., CBASS runs 1 and 2), we kept surviving F003 and H2 anemones from the consecutive CBASS runs under rearing conditions for one month, after which the surviving anemones were subjected to a third CBASS run (anemones subjected to 39 °C as well as some 34 °C- and 36 °C-treated animals died during the one-month interval). As before, photosynthetic efficiencies were measured 1 hour after ramping down from heat stress and dark acclimation (Figure 2 A, PAM 3). Notably, temperature tolerance thresholds of surviving anemones were higher after one month (CBASS run 3) than after the second consecutive CBASS run (CBASS run 2). However, population ED50 thermal tolerance thresholds were still lower compared to the initial measurements (CBASS run 1). Thus, anemones exhibited clear signs of temperature tolerance loss following instantaneous repeat exposure to heat stress (short-term). Further to that, we observed lowered thermal thresholds after a longer term period under ambient conditions, although with signs of partial recovery. This suggests the presence of legacy effects arising from prior heat stress exposure that may need to be considered in field-based measurements (Evensen et al., 2022).

### CBASS assays resolve thermal tolerance differences following microbiome manipulation

Microbiome manipulation, e.g., the provision of beneficial microorganisms for corals (BMCs) (Peixoto et al., 2017), is currently discussed as a type of active intervention to increase the resilience of coral (Santoro et al., 2021; Voolstra et al., 2021a). However, standardized phenotype diagnostics that can measure, for instance, a change in bleaching susceptibility following microbial exposure are missing. As a proof-of-principle that short-term heat stress assays can resolve thermal tolerance differences in Aiptasia following microbiome manipulations, we exposed five F003 and H2 anemones each to 10^7^ cells/mL of the coral bleaching pathogen *Vibrio coralliilyticus* 24 hours prior to as well as during a CBASS assay (Figure 3 A). We found that ED50 thermal tolerance thresholds drastically decreased by approximately 3 °C after *V. coralliilyticus* inoculation compared to untreated (control) anemones (F003 ED50_Control_ = 38.07 °C vs. ED50_*Vibrio*_ = 35.69 °C; H2 ED50_Control_ = 37.21 °C vs. ED50_*Vibrio*_ = 34.59 °C) (Figure 3 B, Table S1). Already under medium (34 °C) and high (36 °C) heat stress, F003 and H2 anemones exposed to *V. coralliilyticus* performed much worse than their control counterparts, as evidenced by a much steeper decline of the photosynthetic efficiency-based ED50 log-logistic regression curves (Figure 3 C). Notably, the response to heat stress following bacterial inoculation was more variable than the response to heat stress alone. This is shown by higher standard errors of population ED50s for F003 and H2 anemones, probably due to the additional source of stress that may affect different anemones to varying degrees (Figure 3 B).

**Figure 3.**
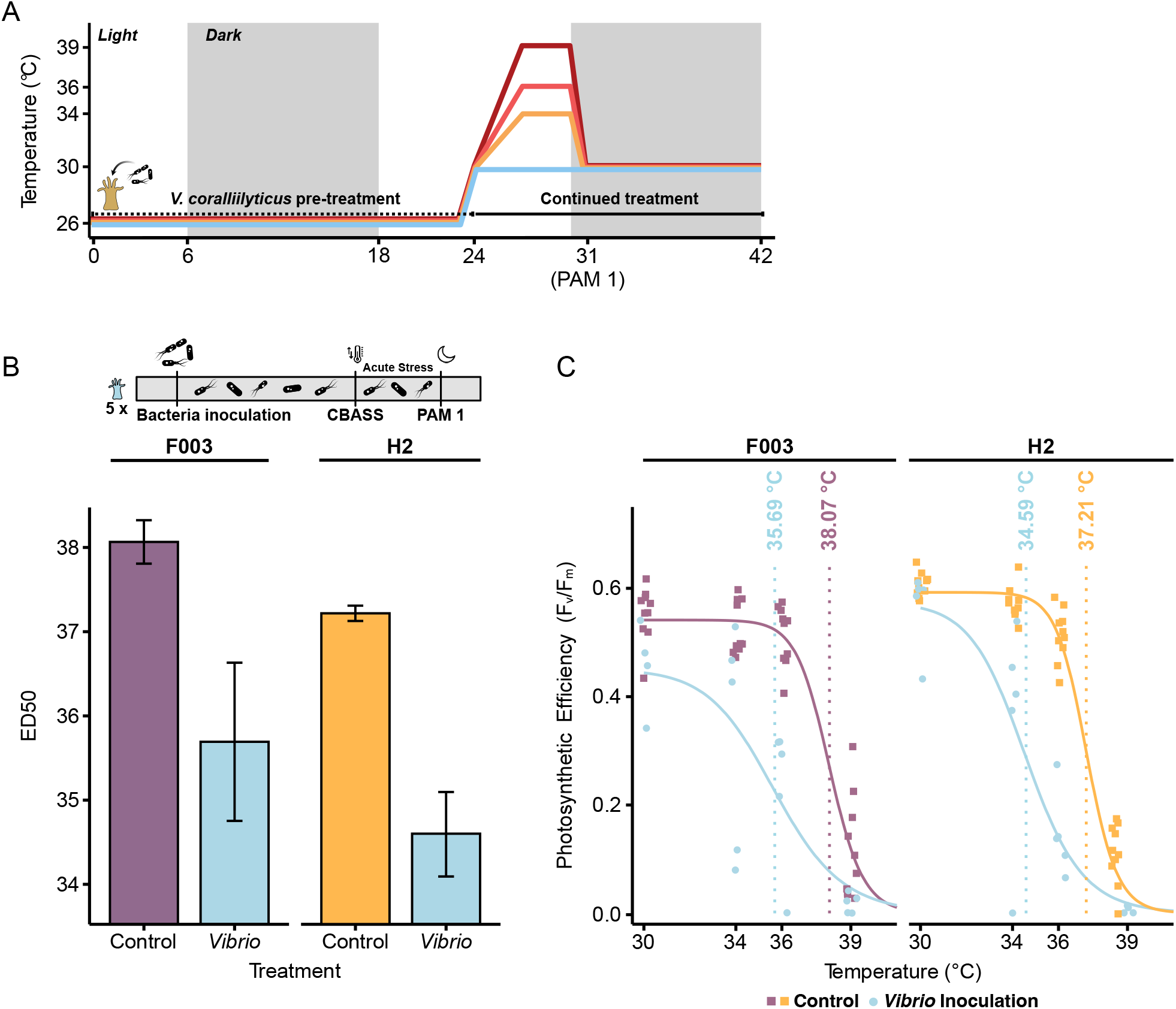
CBASS-based thermal tolerance thresholds decrease following incubation with a coral bleaching pathogen. **(A)** Schematic representation of microbiome manipulation through bacterial inoculation with *V. coralliilyticus* and subsequent CBASS testing. Inoculation was started 24 h prior to the CBASS assay and continued for the duration of the run. The PAM fluorometry measurement time point and CBASS profile are indicated on the x-axis. **(B)** Population ED50 thermal tolerance thresholds based on photosynthetic efficiency (F_v_/F_m_) measures of untreated control and bacteria-inoculated F003 and H2 anemones. Population ED50 thermal tolerance thresholds of both Aiptasia strains decrease drastically after heat stress subsequent to *Vibrio* inoculation. Error bars represent the standard error of the population ED50s (Control, n = 10 anemones per temperature profile; *Vibrio*, n = 5 anemones per temperature profile). **(C)** Photosynthetic efficiencies (F_v_/F_m_) of untreated control and *Vibrio*-inoculated anemones across CBASS assays of F003 and H2 anemones. Population ED50 thermal tolerance thresholds based on log-logistic regression curves of F_v_/F_m_ measurements are denoted as vertical lines.

## Discussion

This study aimed to assess the suitability of short-term acute heat stress assays using CBASS as a standardized screening platform to assay phenotypic effects of microbiome manipulations on thermal tolerance for Aiptasia, a model system for coral-microbe symbiosis (Röthig et al., 2016; Costa et al., 2021). Our results demonstrate that thermotolerance phenotypes are highly reproducible for two strains of Aiptasia, F003 and H2 (Figure 1). We could further show that subsequent heat stress exposures affect thermal tolerance thresholds and that even after a month-long rearing under ambient conditions, legacy effects in the form of reduced thermal thresholds are measurable, although with some signs of recovery (as shown by the regained higher thermal thresholds following month-long rearing in comparison to the loss of thermal tolerance following the two consecutive thermal stress assays) (Figure 2). Lastly, we could show that CBASS assays resolved thermotolerance differences following inoculation with the coral bleaching pathogen *V. coralliilyticus* (Ben-Haim and Rosenberg, 2002; Gibbin et al., 2019) (Figure 3), furthering the notion of CBASS as a suitable tool to assess effects of bacteria on thermal tolerance.

### Aiptasia exhibit strain-specific thermal tolerances

F003 anemones showed overall higher ED50 thermal tolerance thresholds, although their heat stress response was more variable than that of H2 anemones. Our observation of these strain-specific differences may be affected by differences in Symbiodiniaceae community composition of the two Aiptasia strains. Whereas H2 exclusively host *Breviolum minutum* (SSB01) algae, F003 associates with a mixture of SSB01, a variant of SSB01, and *Symbiodinium linucheae* (SSA01) (Xiang et al., 2013; Grawunder et al., 2015). Since SSB01 are generally thought to be more tolerant to high temperatures than SSA01 (Russnak et al., 2021), varying algal community compositions of F003 anemones might affect their physiological response leading to more variable thermal stress responses. Besides algal association, anemone size could also have a role, as F003 anemones were overall slightly bigger than H2 anemones. Further, F003 anemones were more variable in size, which may contribute to the higher segregating variance of population ED50s observed in the repeat CBASS run. Following the same rationale, the population of H2 anemones was more uniform in size, and population ED50s were less variant.

One experimental aspect that we consider critical is the timing of PAM fluorometry measurements. As our repeat CBASS cycling experiment demonstrates, population ED50s change considerably over time as a consequence of heat stress exposure. Thus, F_v_/F_m_ measures at precisely one hour after the temperature ramp down and dark acclimation are central to successfully standardizing ED50 thermal tolerance thresholds. We also found that measuring photosynthetic efficiency at this time point provides the least amount of variance and most reproducible results for both strains, in comparison to the next morning overnight recovery time point (Figure S4). Moreover, measurements 1 hour after the heat-hold save time and resources, as thermal tolerance thresholds can be determined within a day.

### The effect of repeat heat stress exposures on thermal tolerance thresholds

Our assessment of anemone thermal tolerance following repeated thermal stress cycling showed decreasing thermal tolerance thresholds after two subsequent CBASS runs. Thermal tolerance thresholds, however, slightly recovered after month-long rearing under ambient conditions. Thus, it is unclear at present whether an extended recovery period would return thermal tolerance thresholds back to their initial levels or even increase them (acclimation). However, our observation of sustained lowered population ED50 thermal tolerance thresholds one month after the initial thermal stress exposures (CBASS run 1 and 2) rather suggests a negative legacy effect of prior heat stress, as observed previously (Grottoli et al., 2014; Hughes et al., 2017; Santoro et al., 2021). The notion of heat stress carry-on effects following thermal stress has consequences for our interpretation of thermal tolerance thresholds from field-based coral measurements. As noted previously (Evensen et al., 2022), corals along the Red Sea gradient exhibited increased thermal thresholds in accordance with higher mean summer maximum temperatures, except for the southernmost site that had experienced a recent bleaching event. Thus, while CBASS assays are highly reproducible, projected thermal tolerance thresholds are affected by prior thermal stress events. It also remains to be determined whether non-stressful seasonal temperature fluctuations can also affect ED50-based thermal thresholds. Given the uncertainty associated with the time it takes for coral and Aiptasia to exhibit unaffected thermal tolerance, acclimate, or to what extent the level of thermal stress affects subsequent recovery, CBASS could help elucidate the time scale for a complete recovery from heat stress.

### CBASS as a functional screening tool for bacteria-host interactions

The contribution of bacteria to host biology and ecosystems is widely acknowledged (McFall-Ngai et al., 2013). This led to the proposal of active interventions surrounding the manipulation or monitoring of microbes to restore ecosystem and animal health (Voolstra et al., 2021a; Peixoto et al., 2022). However, we are still missing a mechanistic and functional understanding at large of how microbes exert an effect on their putative hosts, e.g. increase stress resilience (Santoro et al., 2021), or what microbes are suitable targets for probiotic intervention (Schultz et al., 2022). An important aspect facilitating the study of the effects of bacteria on host/holobiont phenotypes is a standardized experimental framework that can produce diagnostic phenotypes. This in turn can then be aligned to subsequent molecular investigations to generate hypotheses as to the underlying molecular mechanisms governing observed differences. Here we demonstrate that anemone thermal tolerance thresholds decrease significantly following inoculation of *V. coralliilyticus. V. coralliilyticus* is a known coral pathogen reported to cause bleaching and necrotic tissue lysis, in particular at temperatures above 27 °C (Ben-Haim and Rosenberg, 2002). This is in line with our observation that anemones under control conditions of 30 °C already showed reduced photosynthetic efficiencies. The concentration of *V. coralliilyticus* used in our setup was high (10^7^ cells/mL) in comparison to what was used in a previous study (10^5^ cells/mL directly applied on the coral fragment, then placed into 1.3 L seawater tanks) (Rosado et al., 2019). However, we did not observe a decrease in thermal tolerance using lower concentrations of *V. coralliilyticus* (concentrations used: 10^4^ cells/mL and 10^5^ cells/mL, data not shown). This might be due to the short-term nature of CBASS assays, which require that measurable effects need to manifest within hours, as previously noted (Alderdice et al., 2022b). Nevertheless, pre-incubation with candidate bacteria prior to CBASS assaying might partially ameliorate this limitation. Thus, we expect that longer pre-incubation times at lower concentrations can substitute for higher bacterial loads.

Using *V. coralliilyticus* inoculation as a proof-of-concept, we demonstrate that CBASS could resolve Aiptasia phenotype differences following microbiome manipulation. Thus far, we have shown the measurable negative effects consequential to pathogen inoculation. In the future, microbiome testing will be conducted in reference to inoculation with a control bacterial strain (e.g., *E. coli*) to rule out non-specificity of effects as a result of bacterial load. In the same vein, although we expect positive effects of beneficial microorganisms to be resolved, the ability of CBASS assays to detect increases in thermal tolerance following bacterial incubation, as would be the case in a screening application for putative probiotic taxa, remains yet to be shown.

Given that the ability to enhance the coral microbiome towards coral stress tolerance improvement relies on the successful screening and identification of beneficial bacteria, we show that the CBASS platform provides an efficient and standardized screening tool that can be used in conjunction with microbiome manipulation efforts and subsequent molecular investigations. The rapidity of the CBASS system in combination with employing Aiptasia as a coral model offers the opportunity for screening a large number of bacteria to identify putative beneficial microorganisms for corals (pBMCs) and elucidate their mechanistic underpinnings.

## Author Contributions

MD and CRV conceived and designed the experiments. MD, JD, CSM, JVK, HM, and CRV generated data. MD analyzed data with contributions from JD and CSM. MD and CRV interpreted data. MD and CRV wrote the manuscript. CRV provided tools and funding. All authors discussed results and commented on the manuscript. All authors approved the final version of the manuscript.

## Acknowledgments

This study was supported by funding from the University of Konstanz, AFF project “Microbiology of host resilience (MORE): towards a functional understanding of the role of bacteria in stress tolerance”. We thank Annika Guse and lab for providing Aiptasia lines.

## Data accessibility

All PAM data, HOBO logger files, and R scripts for computation of ED50 temperature tolerance thresholds are available in the associated GitHub repository CBASS_ReprodRepeatLegacy_Aiptasia at https://github.com/reefgenomics/CBASS_ReprodRepeatLegacy_Aiptasia

