## Supplementary Material for "Short-term heat stress assays provide a standardized screening tool to assess impacts of microbiome manipulation on Aiptasia thermal stress tolerance"

Affiliation:
Department of Biology, University of Konstanz, Konstanz, Germany.

†These authors contributed equally to this work.

**Table S1 Population ED50 thermal tolerance thresholds of F003 and H2 anemones.** Population ED50 thermal tolerance thresholds are based on photosynthetic efficiency ($F_v/F_m$) measures. Listed are ED50s for the reproducibility experiment across two CBASS systems (PAM 1 CBASS system A and PAM 1 CBASS system B), the repeated heat stress experiment across five time points (PAM 1 - PAM 5), and the pathogen inoculation experiment across control (PAM 1) and *Vibrio*-inoculated (PAM 1 *Vibrio*) anemones.

| Experiment | Strain | CBASS System | Time point | ED50 (°C) |
|---|---|---|---|---|
| ED50 reproducibility | F003 | A | PAM 1 | 37.94 |
| | F003 | B | PAM 1 | 38.07 |
| | H2 | A | PAM 1 | 37.19 |
| | H2 | B | PAM 1 | 37.21 |
| Repeated heat stress | F003 | B | PAM 1 | 38.07 |
| | F003 | B | PAM 2 | 36.43 |
| | F003 | B | PAM 3 | 36.30 |
| | F003 | B | PAM 4 | 36.47 |
| | F003 | B | PAM 5 | 36.95 |
| | H2 | B | PAM 1 | 37.21 |
| | H2 | B | PAM 2 | 36.64 |
| | H2 | B | PAM 3 | 35.60 |
| | H2 | B | PAM 4 | 35.20 |
| | H2 | B | PAM 5 | 36.50 |
| Pathogen inoculation | F003 | B | PAM 1 | 38.07 |
| | F003 | B | PAM 1 *Vibrio* | 35.69 |
| | H2 | B | PAM 1 | 37.21 |
| | H2 | B | PAM 1 *Vibrio* | 34.59 |

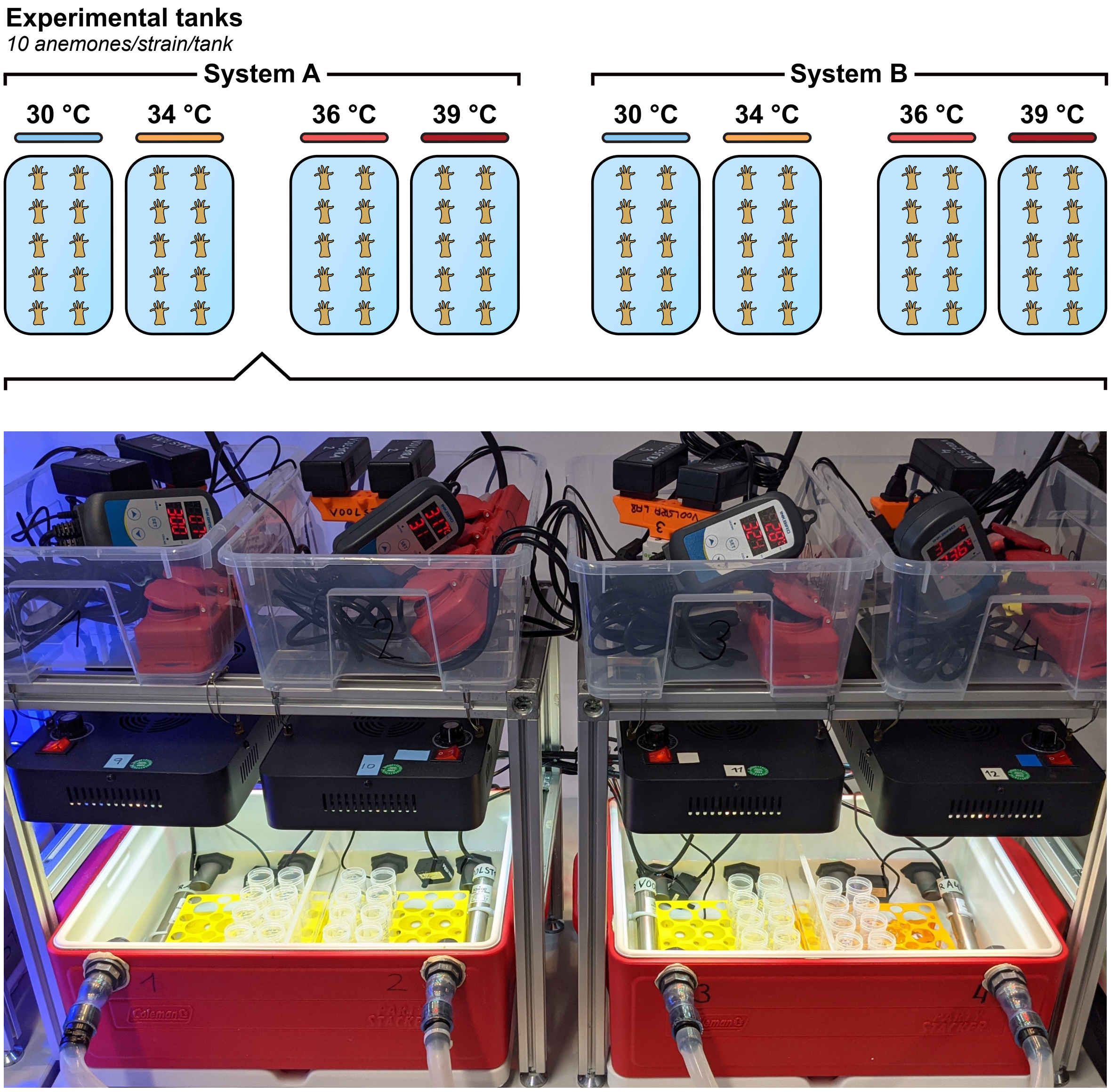


**Figure S1 CBASS experimental setup.** For each anemone strain (F003 and H2), two independent CBASS systems (system A and system B) with four temperature tanks (30 °C, 34 °C, 36 °C, and 39 °C; 10 anemones per tank) each were used side-by-side to determine the reproducibility of thermal tolerance thresholds. Anemones were kept in 25 mL Eppendorf tubes filled with 20 mL FASW. For larger experimental setups, up to 20 anemones fit in each temperature tank, allowing 160 anemones to be tested at the same time. Shown is one CBASS system with four temperature tanks.

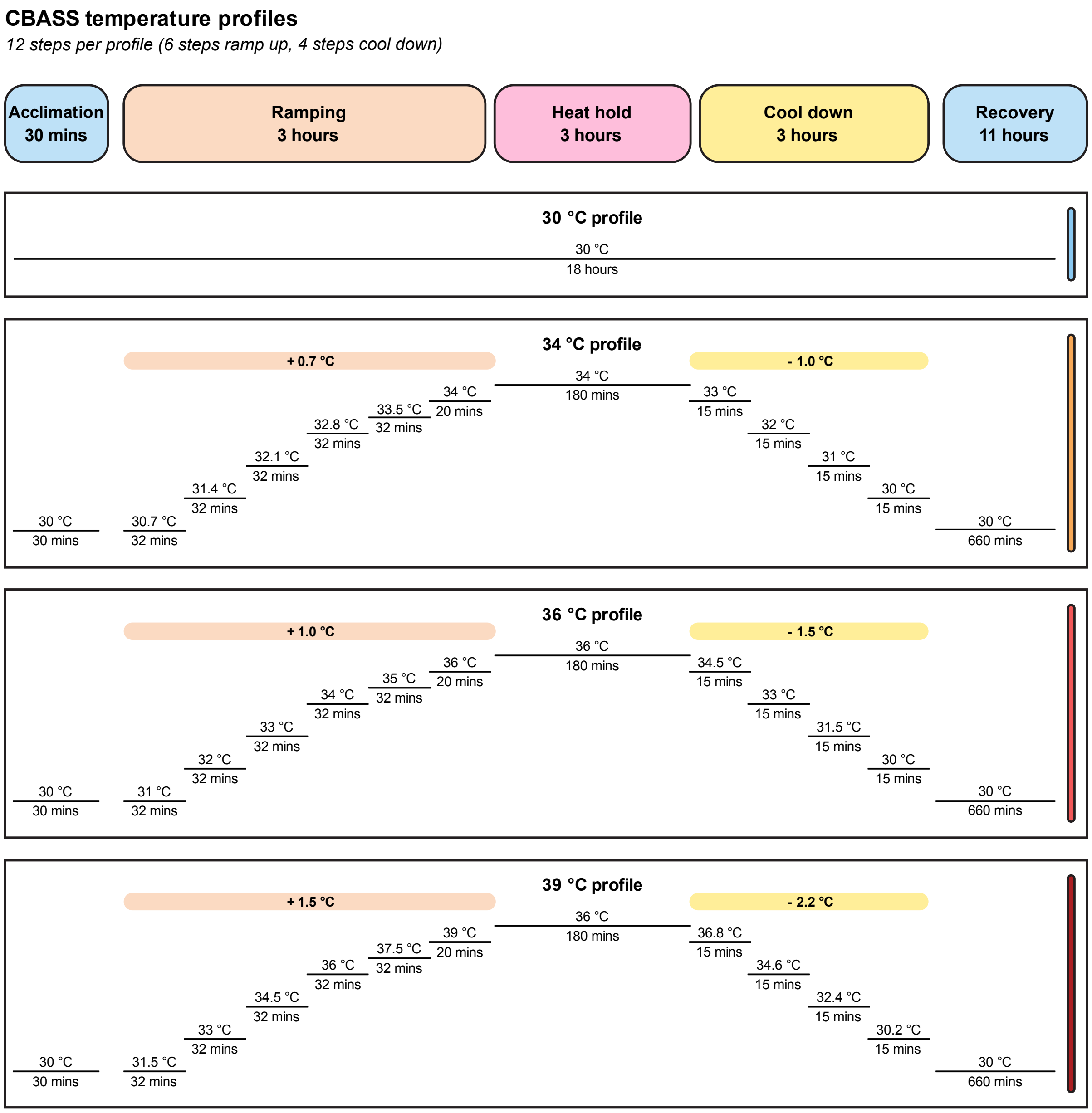


**Figure S2 CBASS temperature profiles programmed on an Inkbird ITC-310T-B.** Prior to heat stress, anemones were acclimated to the 30 °C baseline temperature for 30 min. Heat stress temperatures were then incrementally ramped up in six steps to 34 °C (control +4 °C, medium), 36 °C (control +6 °C, high) and 39 °C (control +9 °C, extreme) over the course of 3 hours. Heat stress temperatures were held for 3 hours and were then cooled down to the baseline temperature (30 °C) in four 15 min-steps. The temperature was kept at 30 °C for an 11-hour (660 min) overnight recovery period.

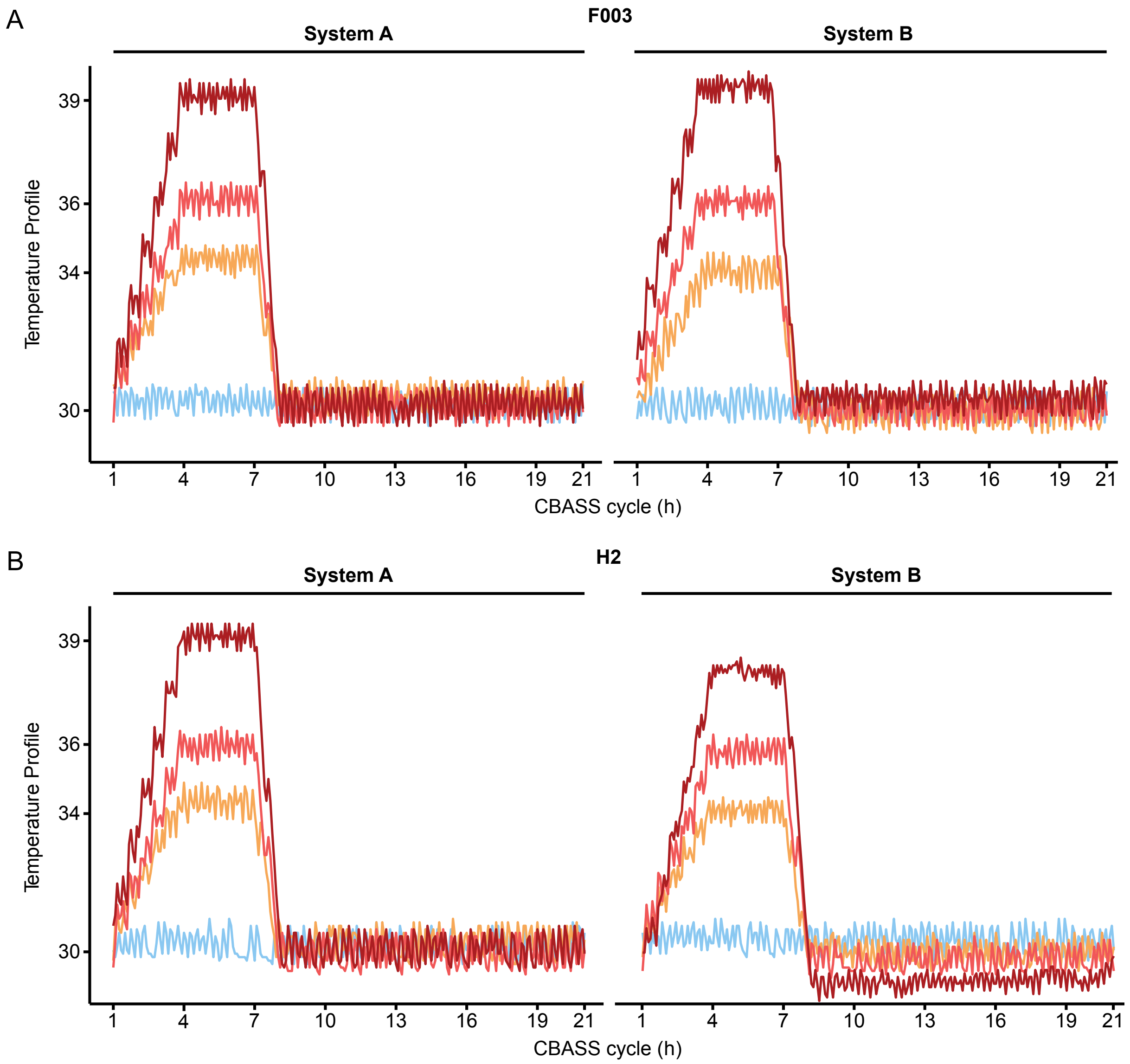


**Figure S3 CBASS temperature profiles for each strain (F003 and H2) and system (System A and System B).** HOBO Pendant Temperature Loggers were used to record the temperature profile accuracy of each treatment tank in 5 min intervals. The control tank temperature profile (30 °C) is shown in light blue, the medium temperature tank profile (34 °C) in orange, the high temperature tank profile (36 °C) in light red, and the extreme temperature tank profile (39 °C) in dark red. Note that the "extreme" temperature tank profile for strain H2 in system B was offset by 1 °C for the reproducibility experiment (CBASS 1 and CBASS 2). Thus, the log-logistic regression to calculate the ED50 thermal tolerance threshold was calculated using 38 °C for the highest temperature and read-out $F_v/F_m$ measurements.

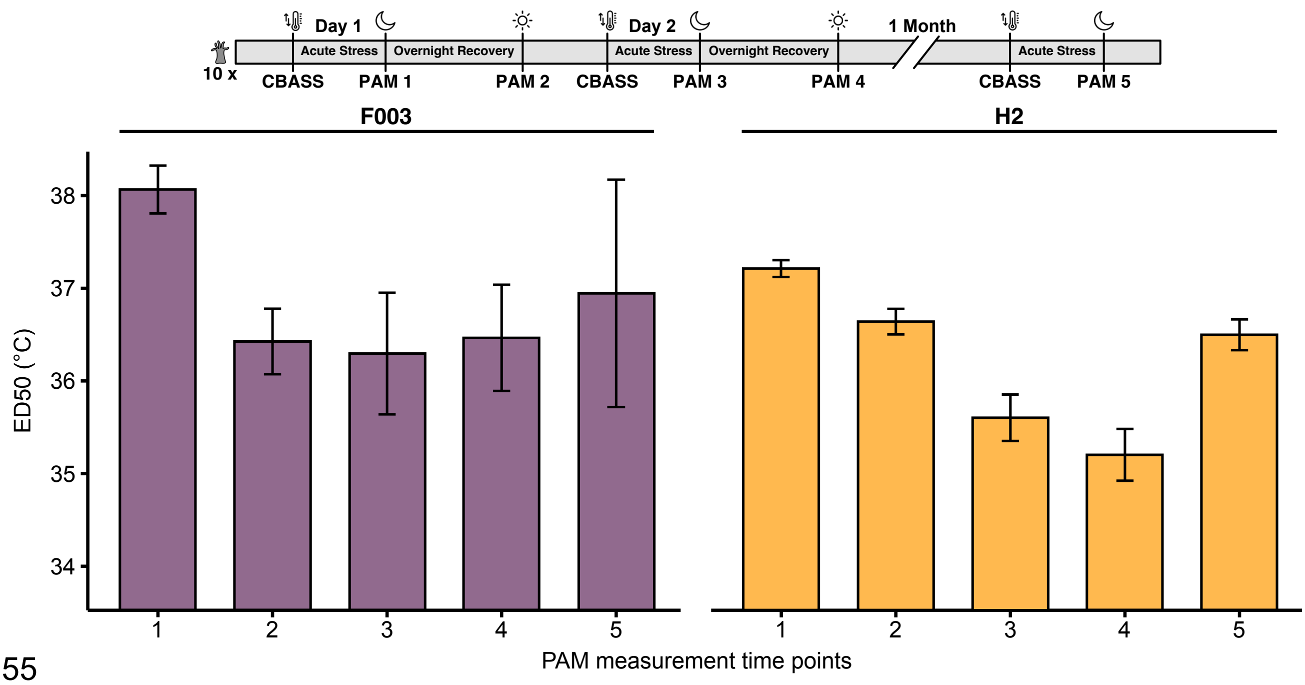


**Figure S4 Comparison of PAM measurement time points and their effect on thermal tolerance threshold determination.** Population ED50 thermal tolerance thresholds were determined for five time points based on photosynthetic efficiency ($F_v/F_m$) measures of Aiptasia strains F003 and H2. Time points one and three are the after heat-hold/ramping down PAM fluorometry measurements, while time points two and four represent PAM fluorometry measurements after an 11 h overnight recovery period. Time point five represents the after heat-hold/ramping down PAM fluorometry measurement of surviving animals after one month under rearing conditions. PAM measurements following heat stress and cooling down (PAM 1) are the least variable and produce the most reproducible results for both Aiptasia strains. Error bars represent the standard error of the respective population ED50.
